# Auditory cortex encodes lipreading information through spatially distributed activity

**DOI:** 10.1101/2022.11.11.516209

**Authors:** Ganesan Karthik, Cody Zhewei Cao, Michael I. Demidenko, Andrew Jahn, William C. Stacey, Vibhangini S. Wasade, David Brang

## Abstract

Watching a speaker’s face improves speech perception accuracy. These benefits are owed, in part, to implicit lipreading abilities present in the general population. While it is established that lipreading can alter the perception of a heard word, it is unknown how information that is extracted from lipread words is transformed into a neural code that the auditory system can use. One influential, but untested, hypothesis is that visual speech modulates the population coded representations of phonetic and phonemic features in the auditory system. This model is largely supported by data showing that silent lipreading evokes activity in auditory cortex, but these activations could alternatively reflect general effects of arousal or attention, or the encoding of non-linguistic features such as visual timing information. This gap limits our understanding of how vision supports speech perception processes. To test the hypothesis that the auditory system encodes visual speech information, we acquired fMRI data from healthy adults and intracranial recordings from electrodes implanted in patients with epilepsy during auditory and visual speech perception tasks. Across both methods, linear classifiers successfully decoded the identity of silently lipread words using the spatial pattern of auditory cortex responses. Examining the time-course of classification using intracranial recordings, lipread words were classified at significantly earlier time-points relative to heard words, suggesting a predictive mechanism for facilitating speech. These results support a model in which the auditory system combines the joint neural distributions evoked by heard and lipread words to generate a more precise estimate of what was said.

**Significance Statement:** When we listen to someone speak in a noisy environment, watching their face can help us understand them better, largely due to automatic lipreading abilities. However, it unknown how lipreading information is transformed into a neural code that the auditory system can use. We used fMRI and intracranial recordings in patients to study how the brain processes silently lipread words and found that the auditory system encodes the identity of lipread words through spatially distributed activity. These results suggest that the auditory system combines information from both lipreading and hearing to generate more precise estimates of what is said, potentially by both activating the corresponding representation of the heard word and suppressing incorrect phonemic representations.

## Introduction

Visual speech improves auditory speech perception during face-to-face conversations (1, 2). These benefits are strongest in noisy situations (3) and in individuals with hearing loss due to healthy aging (4), intrinsic brain tumor (5), stroke (6, 7), concussion (8, 9), or cochlear implants (10). However, there is limited understanding of how the brain enables vision to facilitate hearing processes.

The ability to extract useful information from visual speech signals (e.g., lipreading) is an implicit behavior that is rooted in the statistical relationship between auditory and visual cues in the natural environment (11). Lip dynamics are strongly correlated with different features of speech including temporal information (onset of words, rate of speech, and the boundaries between words) and relative spectral pitch based on the acoustics of the oral cavity (1). Most recognizably, the shape of the lips during speech is reliably associated with corresponding speech sounds (12); these simple lip shapes are described as visemes and are analogous to phonemes in the auditory domain (basic units of speech sounds).

Research has demonstrated that silent visual speech (e.g., lipreading) evokes activity within the auditory system (13, 14). Indeed, intracranial electroencephalography (iEEG) recordings indicate that visual speech influences processing in auditory regions through multiple temporal, spectral, and spatial configurations (15). While these findings highlight the broad influence of visual information on auditory speech processing, differences in activity do not provide a mechanistic account for how visual speech signals integrate with auditory neuronal populations. Among potential mechanisms, it is best understood that visual timing information during continuous speech biases auditory timing through phase-resetting mechanisms (16, 17). However, it remains unclear how lipreading information (visemes) is transformed into a signal used by the auditory system.

Within the auditory domain, phonetic and phonemic features are encoded by local and distributed populations of neurons, respectively (18, 19). Mesgarani and colleagues (18) used human iEEG recorded from high-density electrodes to demonstrate that phonemes are represented by distributed populations of neurons in the STG. Combined with past research, these data support a model in which the STG contains a patchy distribution of neurons that are tuned to specific phonetic features via their spectro-temporal profiles (18, 20). For example, data show spatially distinct responses in these regions to spectrally similar phonemes such as /ba/ and /da/ (20–22), and clustered responses across a large phoneme-space (e.g., the distributed pattern of activity to /ma/ is more similar to /na/ than it is to the spectro-temporally distinct phoneme /ba/ (18)). Indeed, the identities of different *heard* phonemes can be decoded by the distribution of activity in the auditory cortex (23), even when the physical auditory stimulus remains the same.

Building on this framework of auditory perception, we and others proposed that activity from lipread visemes is relayed from visual regions to auditory cortex, preferentially modulating the same populations of neurons that encode matching phoneme responses (15, 24). In this hypothesis, heard and lipread activations in auditory cortex are combined through a winner-take-all mechanism, in which the phoneme population with highest activation profile leads to the phoneme that is perceived (24).

Here we test the hypothesis that the identities of individual visemes are represented in the auditory system through distributed patterns of activation, and these spatial distributions match corresponding phoneme representations. Auditory cortex activation magnitude and informational content were examined using functional magnetic resonance imaging (fMRI) in healthy individuals and iEEG recordings in patients with epilepsy during word perception tasks, in which patients either saw the lip movements or heard the speech sounds for the same groups of words. The identities of the different words were classified from fMRI and iEEG signals in auditory cortex using support vector machine classifiers (SVMs). Results demonstrate that the auditory system reliably encodes the identity of visemes using spatially distributed activity in a similar manner to heard words. Moreover, visemes evoked spatially similar activity to matching phonemes, consistent with the hypothesis that visual speech targets corresponding phoneme representations.

## Results

### fMRI Experiment

In Experiment 1 we presented subjects (*n* = 64) with consonant vowel (CV) syllables while fMRI activity was acquired. Each trial included the silent video or auditory stimulus taken from a speaker producing the CVs /mama/, /fafa/, or /kaka/ (Fig. 1). Stimuli were presented using an optimized event-related design and data were analyzed using univariate and decoding approaches to examine the activation and information present in heard and lipread signals. Simultaneous auditory-visual stimuli were not included in this design to ensure adequate signal-to-noise ratios for classifying silent visual-only speech.

**Figure 1.**
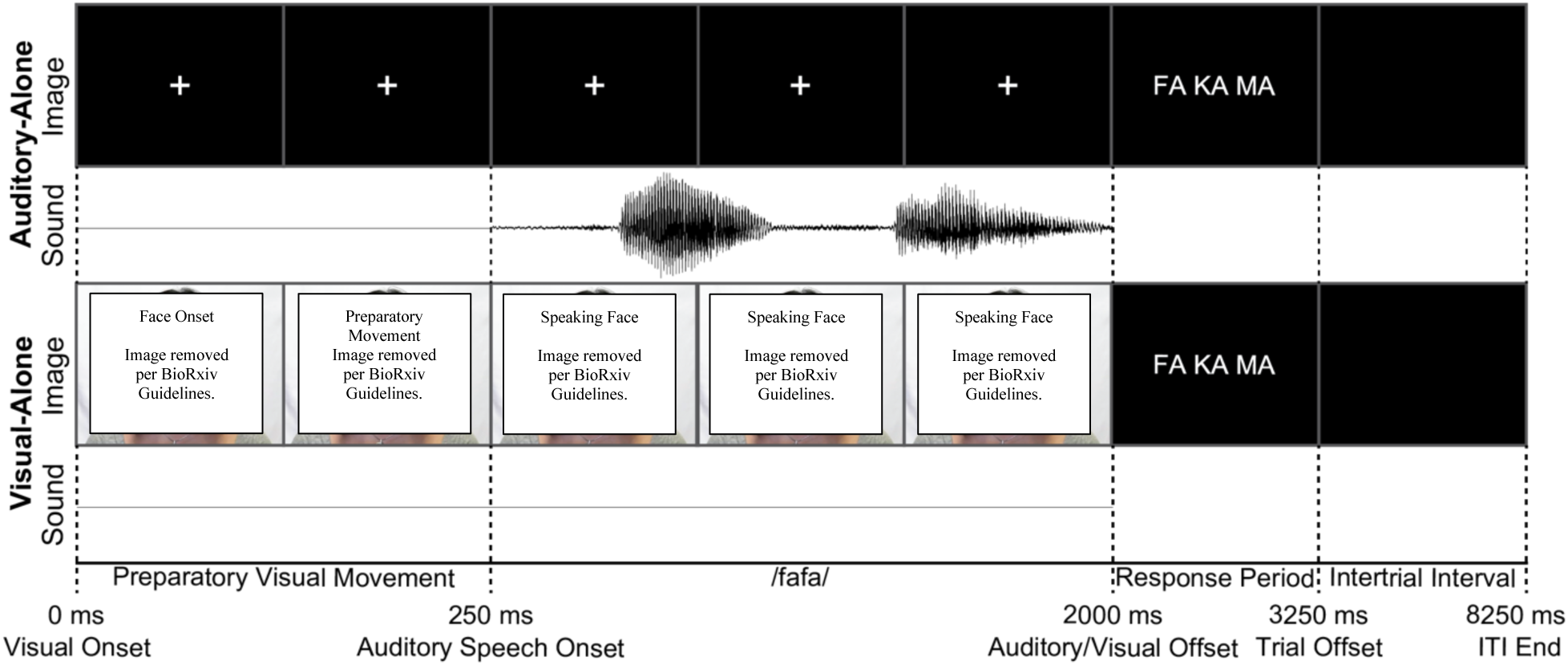
fMRI task schematic. Schematic of auditory and visual trials. Auditory trials began with a fixation cross followed by a CV stimulus (either /mama/, /fafa/, or /kaka/). Visual trials presented the visual components of these same recordings without the corresponding audio track. After stimulus offset, subjects were cued to identify which of the three phonemes (or visemes) they saw (or heard) via button press.

Mean behavioral accuracy was high in both conditions: 95.67% (SD = 3.01%) in the auditory-only condition and 92.31% (SD = 3.72%) in the visual-only condition. As expected, the mean accuracy in the auditory-only condition was significantly higher than the visual-only condition; *t*(63) = 6.57, *p* < 0.001, *d* = 0.96. None of the 64 subjects performed below the pre-registered exclusion threshold (accuracy in either condition below 75%).

Previous research demonstrated that silent lipreading increased fMRI BOLD responses within auditory regions (13). First, we replicated this finding using univariate contrasts in the auditory-only and visual-only conditions (phonemes vs fixation and visemes vs fixation). Fig. 2a-c shows the results of the whole-brain analyses, corrected for multiple comparisons using cluster statistics (vertex-wise threshold of P < 0.001, cluster-corrected to P < 0.05, frontal regions excluded as described in the Methods but see Supp. Fig. 1 for unmasked data). Complete statistics for each analysis are reported in Supp. Tables 1-3 and beta estimates extracted from auditory and visual regions of interest (ROIs) are shown in Supp. Fig. 2. Phonemes elicited significantly increased BOLD activity within the STG bilaterally and decreased BOLD within visual regions. Visemes showed a similar pattern in the auditory system with increased BOLD in the STG and pSTS bilaterally, along with increased BOLD in visual regions, including hMT+, the fusiform gyrus, and visual cortex. Importantly, phonemes elicited stronger activity relative to phonemes throughout the STG.

**Figure 2.**
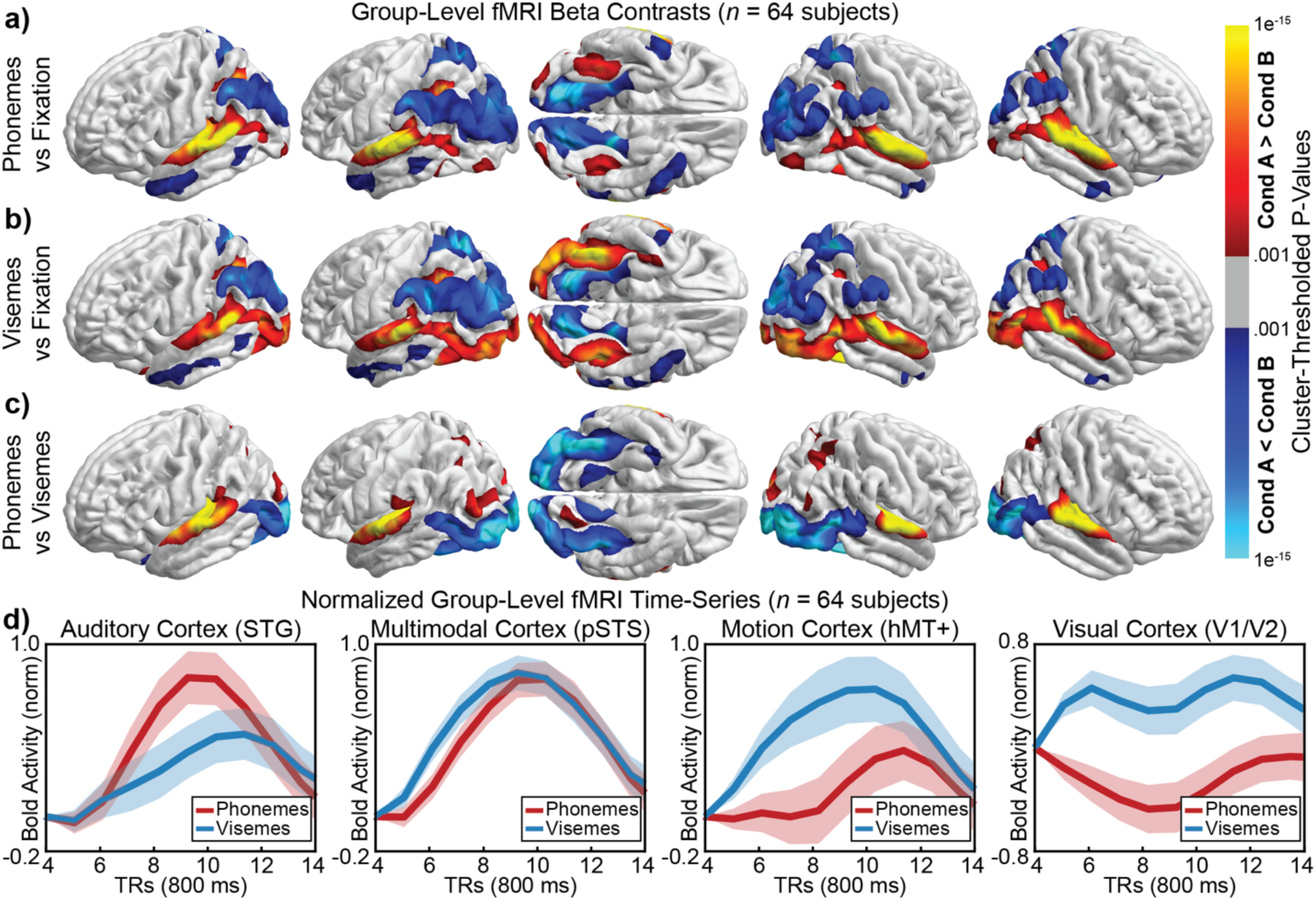
Univariate activations. (a) phonemes vs fixation (b) visemes vs fixation, and (c) phonemes vs visemes. Phonemes evoked maximally increased activity in the STG bilaterally. Visemes evoked increased activity within bilateral visual cortex, left pSTS, right MT, and the STG bilaterally, though significantly weaker within the STG compared to phonemes. Colored regions reflect significant increases (red and yellow) or decreases (blues) in task-related activation (thresholded at p<.001 and corrected for multiple comparisons using cluster-statistics). (d) BOLD time-series extracted from four regions (averaged across left and right hemispheres). Data were normalized based on the maximum average value between 3.2 (TR=5) and 9.6 (TR=13) seconds to better visualize relative shapes of the time-series. Notably, phonemes and visemes showed a similar temporal response in the STG but visemes elicited activity in the pSTS more quickly than phonemes. Stimulus onset occurred at TR=1.

Fig. 2d shows the normalized BOLD time-series from four regions of interest. Notably, BOLD response timing in the STG was similar across phoneme and visemes, consistent with a model in which visual speech biases early auditory perceptual processes. Conversely, visemes elicited activity with the pSTS significantly before phonemes, suggesting a potential pathway for the transformation of visemes to phonemes (consistent with previous studies; e.g. (25)). For example, at 3.2 seconds following stimulus onset, visemes evoked significantly greater activity in the pSTS, *t*(63) = 5.68, *p* < .001 (FDR-corrected), *d* = 0.710. These plots show the time-series scaled to have the same amplitude across conditions to better see similar onset times in the STG and different onset times elsewhere. Unnormalized data are shown in Supp. Fig. 3.

Univariate contrasts reveal activation magnitude but not informational content or representational structure. Previous decoding-based approaches using fMRI (26, 27) and iEEG (28, 29) demonstrated that auditory speech identities could be reconstructed from spatially distributed activity in auditory cortex. To examine whether viseme identity is represented in the auditory system by distributed patterns of activity, we used multivariate pattern analysis (MVPA) (30) to classify individual phoneme and viseme labels.

Whole-brain searchlight-based MVPA was applied at the individual-subject level (31) conducted separately for each of the two conditions of interest (auditory-only and visual-only trials). Results of the whole-brain analysis, corrected for multiple comparisons using cluster statistics are shown in Fig. 3 (vertex-wise threshold of P < 0.001, cluster-corrected to P < 0.05); full statistics for each analysis is reported in Supp. Tables 4 and 5. In the auditory-only condition, peak decoding accuracy was observed bilaterally in the STG and pSTS. This is consistent with previous studies demonstrating phonetic representations in the STG using MVPA (32). In the visual-only condition, peak decoding accuracy was observed within the STG bilaterally, the left pSTS, visual cortex bilaterally, and right hMT+. To understand the spatial overlap of phoneme and viseme representations in the auditory system we compared the spatial distribution of the classification maps. Within the STG, results showed that a majority of vertices contained either only phoneme information or both phoneme and viseme information, with very few vertices representing viseme information alone (Fig. 3c). Across the left and right STG, phonemes (but not visemes) were significantly classified at 27.2% of vertices, phonemes and visemes were jointly classified from 25.2% of vertices, and visemes (but not phonemes) were significantly classified at 0.16% of vertices. In total, phonemes were classified at twice as many vertices compared to visemes within the STG (52.4% vs 25.4%).

**Figure 3.**
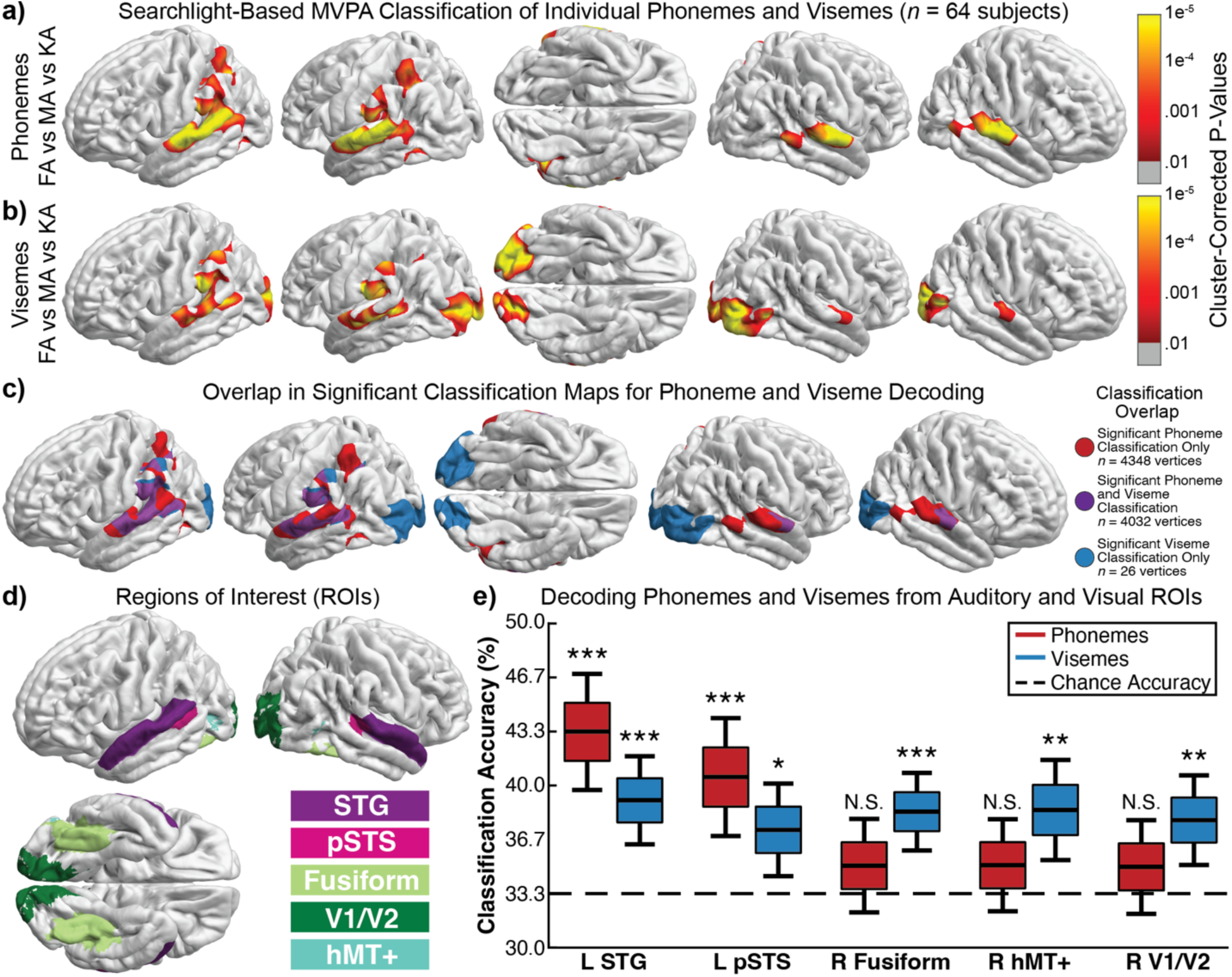
fMRI decoding of phoneme and viseme information in an event-related design. (a-c) Searchlight-based MVPA classification in *n* = 64 subjects. Classifiers were trained to identify (a) the phoneme heard (/fafa/, /mama/, or /kaka/) in the auditory-only condition, (b) the viseme seen in the visual-only condition, or (c) condition differences between auditory-only and visual alone trials. Decoding was conducted at the individual subject level and only group-level differences greater than chance (thresholded at p<.001 and corrected for multiple comparisons using cluster-statistics) are shown. (a) Peak phoneme decoding was observed in the bilateral STG. (b) Significant viseme decoding was observed in the bilateral STG, left pSTS, and visual regions. (c) Vertices with significant classification of phonemes but not visemes (red), visemes but not phonemes (blue), or with significant classification of both phonemes and visemes (purple). There is a large overlap in the vertices at which visemes and phonemes could be classified. Restricted to the just the STG, vertices at which viseme classification was significant covered roughly half of the area in the STG that phonemes were classified successfully at (48.1% overlap) with negligible area uniquely able to classify visemes. (d) Regions of interest (ROIs) used for hypothesis driven classification at the single-subject level. (e) Results of classification at selected ROIs. Phonemes were significantly classified from the left STG and pSTS. Visemes were significantly classified from the left STG (consistent with the hypothesis that information about visemes is represented within the STG), pSTS, and visual cortex. Center line reflects the mean, colored box SE, and the tails 95% confidence intervals. *p<.05, ***p<.001. Chance accuracy is 33.3%.

The univariate analysis showed that visual speech increased BOLD activity broadly across auditory regions and viseme decoding analyses identified above-chance classification accuracy in a more restricted set of vertices throughout the STG. Comparing the two results, the univariate visual-only analysis showed two times as many significant vertices in the STG (bilaterally) compared to the area with significant viseme classification in the MVPA analysis (56.7% vs 25.4% of STG vertices). This is consistent with the prediction that only a restricted proportion of the STG encodes visemic information, while other regions reflect domain general responses to the visual signals or the presentation of other visual information (e.g., temporal or spectral information (1)). Conversely, the auditory-only analysis showed significant univariate responses in 79.2% of the STG, and phonemes were significantly classified from 52.4% of vertices within the STG.

To further quantify the relative information across target regions, we performed individual-subject SVM classification in five regions of interest (ROIs) in each hemisphere (dimension of vertices within the ROI; Fig. 3d) and corrected p-values using FDR. As shown in Fig. 3e and Supp. Fig. 4, phoneme classification accuracy was strongly above chance (33.3%) in the left and right STG and pSTS (all *p* < .001). Viseme classification accuracy was similarly above chance in the left STG (*p* < .001) and left pSTS (*p* < .05). Visemes were additionally able to be classified from multiple visual regions, most notably in right hMT+ (*p* < .001), right fusiform gyrus (*p* < .01), and right V1/V2 (*p* < .01). To confirm that our viseme decoding results were generated from auditory selective regions as opposed to non-selective regions of the STG, we repeated viseme classification only at voxels within the STG that showed significant increase in BOLD activity to phonemes relative to fixation. These auditory selective results showed a similar pattern of effects with significant viseme classification in both the left STG (*p* < .001) and right STG (*p* < .05).

### iEEG Experiment

Results from the fMRI study demonstrated that viseme information is represented in auditory areas. Moreover, because visemes were classified based on the spatial distribution of vertices in the STG, this supports a model in which lipreading is represented through population-coded responses in the auditory system, similar to the neural representation underlying phonemes(18). However, the slow temporal dynamics of fMRI signals prevent a fine-grained analysis of the time-course of lipreading activation to examine when this information is available to the auditory system. Additionally, the use of only three dissimilar CV stimuli prevented a more graded analysis of these population-coded responses, such as whether more perceptually similar phonemes (e.g., /ga/ and /da/) elicit more similar population-coded responses relative to perceptually distinct phonemes (e.g., /fa/ and /ba/). To answer both of these questions we collected data from a similar auditory-visual speech paradigm from *n* = 6 patients with epilepsy who had electrodes implanted within auditory areas of the brain (Fig. 4a). Patients were presented with 240 auditory-only (listening) and 240 visual-only (lipreading) trials containing a single 1-2 syllable word. Each word began with one of four consonants (’B’, ‘F’, ‘G’, or ‘D’) to enable the classification of distinct phonemic patterns. 40 distinct words were used (10 containing each of the 4 initial consonants; Supp. Table 6) and each word was repeated 6 times within each condition. On each trial subjects selected the initial consonant heard or seen from four options (4-alternative forced choice paradigm). Subjects’ mean behavioral accuracy across auditory-only and visual-only trials was significantly above chance (25%) at the group level: auditory-only (M = 91.9%, SD = 8.1%, *t*(5) = 20.2, *p* < .001, *d* = 8.23), visual-only (M = 67.4%, SD = 17.9%, *t*(5) = 5.78, *p* = .002, *d* = 2.36). As expected, auditory-only trials were correctly identified significantly more often than visual-only trials, *t*(5) = 5.93, *p* = .002, *d* = 2.42.

**Figure 4.**
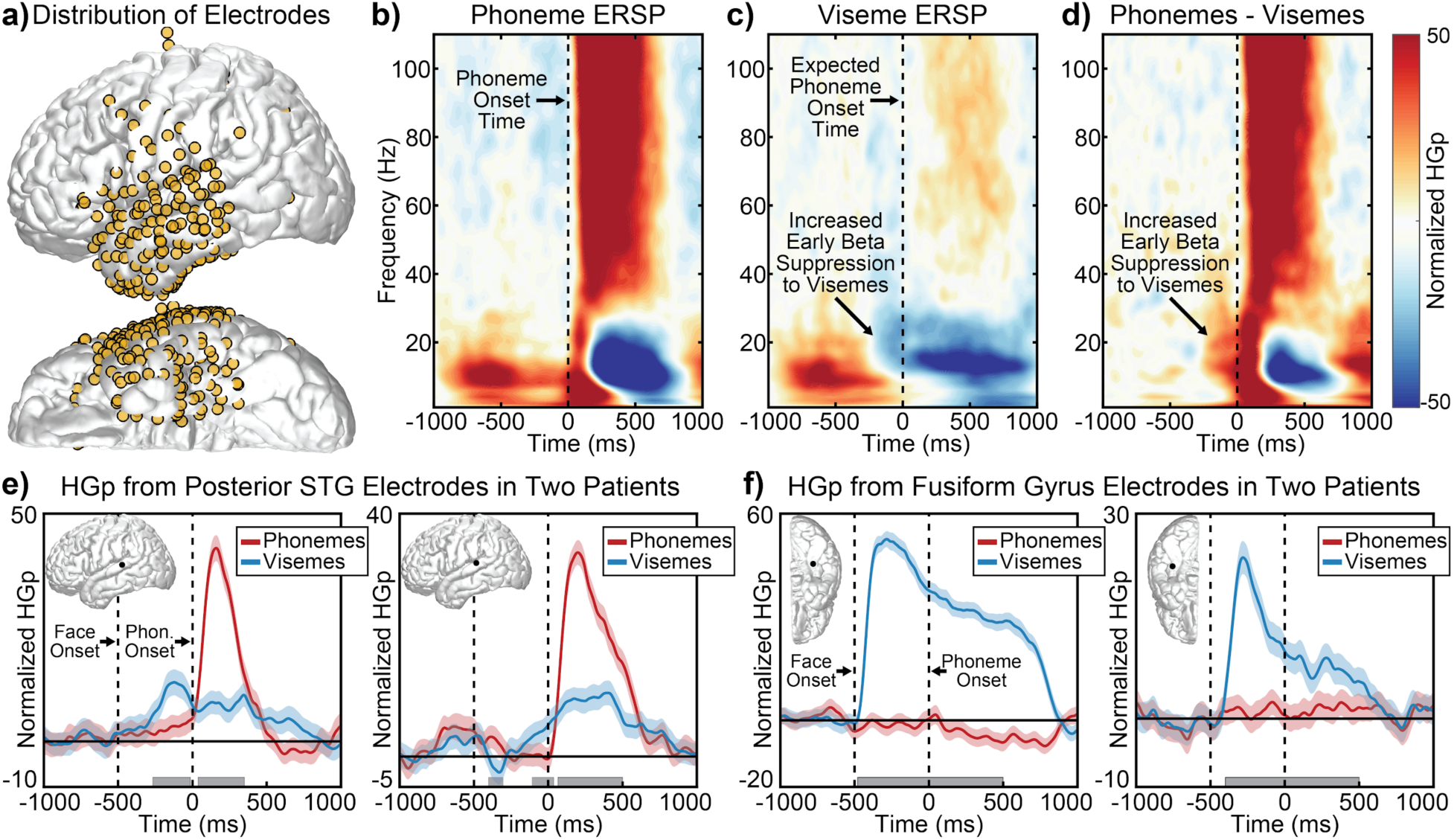
iEEG results during an auditory-only (listening) and visual-only (lipreading) speech perception paradigm. (a) Distribution of all recorded electrodes (those beneath the pial surface not shown) (*n* = 6 patients). (b-d) Event-related spectral perturbations (ERSP) plots from all STG electrodes, averaged across subjects. (b) Phoneme responses peaked after sound onset with theta and high-gamma power (HGp) increases, as well as beta suppression. (c) Viseme responses evoked maximal changes in beta power, with increased beta suppression starting before the expected time of sound onset. (d) Difference between phoneme and viseme ERSP plots. (e) HGp responses from two STG electrodes in response to auditory-only trials (phonemes; red lines) and visual-only trials (visemes; blue lines). Posterior STG electrodes showed increased HGp responses to visemes before the time when speech sounds would be expected to begin. (f) HGp responses from two fusiform gyrus electrodes. Visemes evoked increased HGp shortly after onset of the face, with elevated HGp persisted throughout the visual movement period. Phonemes failed to evoke reliable changes in activity within the fusiform gyrus. Shaded regions reflect single condition 95% confidence intervals. Light gray boxes show significant between condition differences (multiple comparisons corrected using FDR).

Words in both the auditory-only and visual-only conditions evoked activity broadly throughout the STG and MTG consistent with prior work (15). Examining the spectral breakdown of these responses (Fig. 4b-d), phonemes evoked increased theta and high gamma power (HGp) and suppressed beta power following word onset; Supp. Fig. 5 shows the spectral responses for different regions of the STG. In natural speech, visual articulations typically occur before the onset of speech-related sounds (typically within 40 - 200 ms of speech onset (33)). Because of this pre-articulatory visual information, visemes evoked increased beta suppression beginning before the expected phoneme onset time (−250 to 0 ms) [*t*(5) = 2.78, *p* = .039, *d* = 1.13]. This is consistent with past research (15) and indicative of possible lateral(34) or feedback inputs (35–37) into the auditory system.

Additionally, visemes elicited moderate changes in HGp following sound onset, even though no sound was present. Viseme-related HGp increases were maximal at the posterior STG, consistent with past research (15, 38, 39). Fig. 4e shows this pattern in single electrode HGp responses from two patients (both electrodes within the left posterior STG), with HGp changes occurring in response to visemes before phonemes. This pattern was distinct from responses in the fusiform gyrus, at which visemes evoked early HGp increases following the onset of the visual stimulus and no reliable response at any point during auditory-only trials (Fig. 4f).

To replicate the results of our fMRI experiment and examine whether viseme information is represented within auditory regions, we decoded word information using spatial and temporal signals from iEEG electrodes. Fig. 5a shows group-level average classification accuracy for decoding the initial word consonant for auditory-only and visual-only trials. iEEG signals used for decoding analysis were recorded from electrodes from superiortemporal, middletemporal and supramarginal areas during 0-500 ms during each trial and downsampled to 10Hz. Electrodes were only included if they were deemed functionally significant: i.e., regardless of condition they had significant activity using a one-sample t-test (p<0.01, FDR-corrected). Classification was conducted separately in each subject using SVM classifiers on single-trial event-related potential (ERP) responses (60 per consonant-initial auditory and visual words) using time points and electrodes as dimensions. We observed significant classification (evaluated using binomial statistics) in all six patients for both auditory-only (all *p* < .05) and visual-only conditions (all *p* < .05) (single-subject statistics shown in Supplemental Table 7). Similarly, at the group-level we observed classification accuracy reliably above chance for both auditory-only (*t*(5) = 5.39, *p* = .0030, *d* = 2.20), and visual-only trials (*t*(5) = 5.57, *p* = .0026, *d* = 2.27). We additionally observed a trend towards greater classification in auditory-only trials relative to visual-only trials (*t*(5) = 2.08, *p* = .0916, *d* = 0.851).

**Figure 5.**
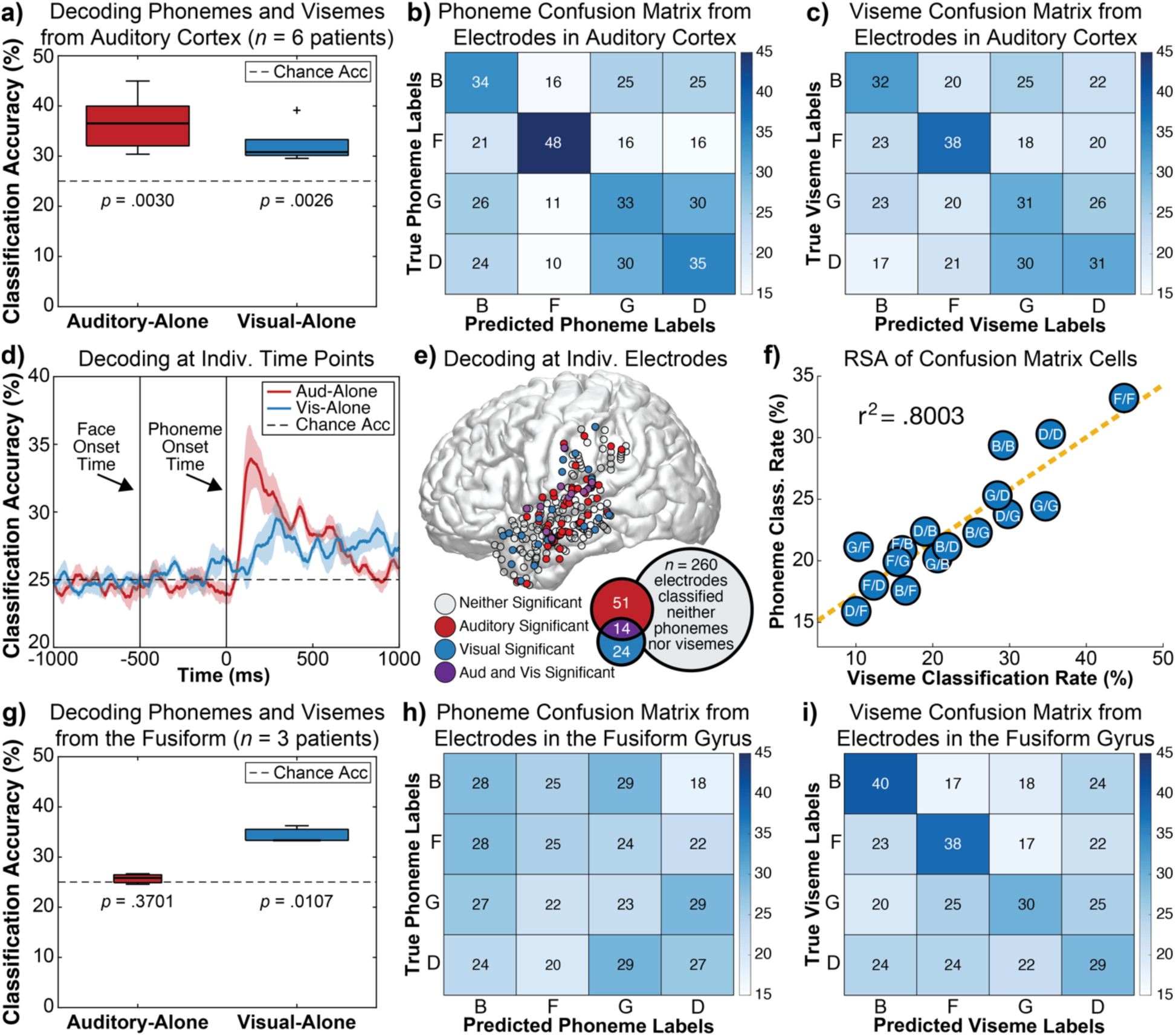
iEEG classification of phoneme and viseme identities from auditory (*n =* 6 patients) and visual (*n =* 3 patients) regions. (a) Accuracy of an SVM classifier in identifying the correct initial consonant (’B’, ‘F’, ‘G’, or ‘D’) from either auditory-only or visual-only words classified at the individual-subject level from spatial and temporal iEEG information. Both visemes and phonemes were reliably classified from auditory electrodes. Chance accuracy is 25% and plots show group-level boxplots. (b-c) Group-averaged confusion matrices taken from 4-class auditory-only and visual-only SVM classifiers. Cells denote the frequency at which each consonant-initial word was predicted (x-axis) relative to the true labels (y-axis). ‘F’ initial words were best classified across both auditory-only and visual-only conditions, whereas ‘G’ and ‘D’ initial words were more readily confused. (d) Group-averaged classification at individual time-points from auditory electrodes (phoneme-onset at 0 sec) showing significant classification accuracy for both auditory-only and visual-only trials shortly after phoneme onset; in the visual-only condition, this time-point reflected the associated speech onset time even though no auditory stimulus was presented. Shaded region reflects SEM. (e) Spatial distribution of electrodes at which auditory-only (red) or visual-only (blue) trials were reliably classified (p<.05 based on binomial statistics); purple electrodes reflect significant classification in both conditions (with 11 out of 14 of these electrodes present in the superior temporal gyrus) and gray electrodes reflect non-significant classification in either condition. Electrodes beneath the pial surface were projected out to the lateral surface for visualization. (f) Scatterplot quantifying the similarity of classification frequency for auditory-only trials and visual-only trials from auditory electrodes (taken from 8-class classifier). Data reflect pairwise classification values, with the first letter reflecting the real consonant label and the second letter the predicted consonant label. For example, ‘F’ trials were predicted correctly at high frequency for both auditory and visual trials, whereas ‘D’ trials were incorrectly labeled as ‘F’ trials infrequently across both auditory and visual trials. The high correlation (r^2^ = .8003, p<.001 permutation test) is consistent with the hypothesis that visual speech evokes responses targeting similar distributions of neurons to corresponding phoneme responses in the STG. (g) Group-level classification accuracy showing that responses in the fusiform gyrus can distinguish between different visemes but not phonemes. (h-i) Group-level confusion matrices for auditory-only and visual-only trials from fusiform gyrus electrodes.

The successful classification of phonemes and visemes indicated that auditory areas represent information about the consonant initial words for both auditory-only and visual-only speech stimuli. The diagonal of the confusion matrices (Fig. 5b-c) shows that this classification was robust for each of the four auditory-only and visual-only stimuli considered (*p* < .05 for 3 out of 4 phonemes and 3 out of 4 visemes) (statistics shown in Supp. Table 8). Previous research has shown that local auditory responses spatially cluster according to phonetic features (18); for the stimuli used here, B, G, and D form one cluster and F another. Consistent with these clusters, classification was higher for auditory words with the consonant F compared to words with the consonants B, G, or D (*t*(5) = 2.95, *p* = .0319, *d* = 1.20); a similar trend was observed for visual-only trials (*t*(5) = 1.80, *p* = .1318, *d* = 0.736), consistent with perceptual ambiguity of these items in phoneme-space.

Following the analyses in the fMRI experiment, we additionally performed a follow-up analysis examining visual-speech information only in electrodes that demonstrated significant auditory activity. Electrodes were identified for inclusion to those within the STG that showed significant auditory responses to phonemes. Even within this more restricted range of electrodes we observed significant viseme classification in four of the six patients at the individual subject level (all p-value less than .01) and significantly greater classification of visemes at the group level relative to chance (*t*(5) = 2.62, *p* = .047, *d* = 1.07).

To examine the time-course of auditory and visual speech representations within the auditory system, we classified the identity of stimuli independently at 1 ms intervals (from −1000 ms to +1000 ms relative to phoneme onset time and smoothed the resultant time-series using a 50 ms moving average). Classification was applied separately for each subject and group-level statistics were calculated across subjects (multiple comparisons were corrected using FDR applied from 0 to 1000 ms) (Fig. 5d). Visemes were reliably classified (all *p* < .05 corrected) at the start of this range from 0 - 23 ms (relative to sound onset) as well as multiple later periods in the time-range. In contrast, phonemes were not classified until 96 ms after sound onset but demonstrated continuous significant classification until 657 ms. Importantly, the early classification of visual-only trials demonstrates that visemic information is available to the auditory system at the same early perceptual stage as is phonemic information, instead of reflecting later categorical or decision-level processes. It is additionally possible that viseme information is available to the auditory system before phoneme information (see ref (40)) but the evidence for this in the current data is weak and requires future confirmation: while classification of visemes was higher than phonemes at 0 ms (*p <* .05 uncorrected), no multiple comparison corrected significant differences were observed between the conditions at any time-point, likely due to the small sample size.

In a parallel set of analyses, we classified the identity of stimuli independently at each electrode within an auditory region (including the STG, MTG, SMG) to understand the spatial distribution of phoneme and viseme classification and their overlap. Phonemes were significantly (*p* < .05 using binomial statistics) classified from 65 out of 260 electrodes (25.0%) while visemes were significantly classified from only 38 electrodes (14.6%). This pattern is similar to the overall classification rate observed in our fMRI data, where phonemes were classified at three times as many vertices compared to visemes within the STG. Restricted to only the STG, 14 electrodes significantly classified visemic information in total, with 11 of these also significantly classifying phonemic information, highlighting the spatial overlap of these processes. Again, this is consistent with the pattern observed in our fMRI data, in which few vertices were sensitive to only visemic information.

Because phonemes are represented through population coded responses, misclassification can reveal information about related neural processes. For example, if the rate at which the consonant /d/ is misclassified as /g/ in both auditory-only and visual-only trials is similar, it suggests similar underlying representations. To test whether auditory cortex shows similar representations for phonemic and visemic information, we calculated a correlation between each of the phoneme-pairs across the phoneme and viseme group-averaged confusion matrices. Fig. 5f shows the scatter plot reflecting classification rate for each consonant pair. Across auditory-only and visual-only trials, classification (and misclassification) rates were highly correlated (*r^2^* = .8003, *p* < .001 permutation test). Significance was calculated by randomly permuting the stimulus labels of each trial and repeating the full classification analysis *n* = 1000 times. This is consistent with our hypothesis that the spatiotemporal neural representation of viseme identities in the auditory areas is similar to that of phonemes.

Three of the six subjects had electrodes along the ventral temporal lobe (including the fusiform gyrus). To examine phoneme and viseme representations in this visual region, we repeated classification on this restricted set of electrodes. Within visual electrodes, group-level classification was significantly above chance for visual-only trials (*t*(2) = 8.74, *p* = .0128, *d* = 5.05) but not auditory-only trials (*t*(2) = 1.15, *p* = .3701, *d* = 0.662). We additionally observed greater classification in visual-only trials relative to auditory-only trials (*t*(2) = 5.29, *p* = .0339, *d* = 3.05). This pattern was seen at the individual-subject level in all three subjects using binomial statistics (all *p* < .01 for visual-only trials and all *p* > .24 for auditory-only trials).

Classification of ERPs revealed robust encoding of phoneme and viseme information in the auditory system, driven by low-frequency oscillatory information (power and phase) that reflects the excitatory/inhibitory balance of local neuronal populations (41). Higher frequency activity (high-gamma power; HGp), in contrast, is associated with the average rate of action potentials generated by a local population of neurons (42). Across HGp from all auditory electrodes, we observed significant classification (evaluated using binomial statistics; *p* < .05) in five out of six patients for auditory-only trials and three out of six patients for visual-only trials. Similarly, at the group-level we observed classification accuracy reliably above chance for both auditory-only (*t*(5) = 3.74, *p* = .013, *d* = 1.53), and visual-only trials (*t*(5) = 3.56, *p* = .0162, *d* = 1.45). We additionally observed a trend towards greater classification in auditory-only trials relative to visual-only trials (*t*(5) = 2.40, *p* = .0614, *d* = 0.981). More reliable classification for low-frequency signals evoked by visemes is consistent with the fMRI finding that classification can occur in auditory regions that do not show increased firing rates in response to visual speech.

## Discussion

Extensive research has shown that silent visual speech can modulate activity within primary auditory regions in humans (13–15, 24, 43, 44). However, multiple sources of information could be contained in these visual-driven auditory responses including visual motion and timing information (45), visual parsing of speech rate (11), visual-derived spectral information (1), general effects on attention or arousal (34), or viseme-to-phoneme transformations (15). Here we tested the hypothesis that the identities of individual visemes are represented in the auditory system through distributed patterns of activation, and these spatial distributions match corresponding phoneme representations. Using fMRI and intracranial electrodes implanted in auditory regions we found that the auditory system reliably encodes the identity of visemes using spatially distributed activity in a similar manner to phonemes. Moreover, visemes evoked similar patterns of confusability as phonemes, consistent with the hypothesis that visual speech activates corresponding phoneme representations.

Data from both fMRI and iEEG showed reliable classification of visemes from auditory regions, maximal in the left pSTS and STG bilaterally. The spatial distribution of significant viseme classification suggest multiple routes from visual cortex (46) to auditory cortex via hMT+ and the pSTS. Consistent with this view, visemes increased BOLD activity in hMT+ and the pSTS before these regions responded to phonemes. Future functional connectivity analyses can be used to examine the paths of transmission of lipreading information from visual to auditory regions and computational analyses to examine how viseme information is transformed into phoneme or phonetically tuned features. For example, dynamic causal modelling (DCM) has previously shown that visual speech modulates auditory processing through ventral and dorsal routes (47).

Classification of iEEG data enables inferences about *when* phoneme and viseme information is available to the auditory system. This temporal resolution is necessary to understand whether visemes are used by the auditory system during the same time window that auditory phonemes are processed in order to support the population-coded perceptual representations of phonemes. Alternatively, if viseme representations follow the perceptual encoding of phonemes (e.g., >500 ms) visemes may instead support categorical decisions about what was heard. The present data showed significant classification accuracy for both auditory-only and visual-only trials shortly after phoneme onset, indicating that visemic information is available to the auditory system at the same perceptual stage as is phonemic information. Nonetheless, it remains possible that silent visual speech can provide visemic information to the auditory system before phoneme onset in cases which visual speech precedes auditory onset (40).

While we observed increased BOLD activity in response to lipreading broadly throughout the STG, this does not necessarily indicate that lipreading increased the spiking rate of excitatory neurons within all auditory regions. Specifically, research indicates that BOLD increases can reflect changes in either local excitation or inhibition as both can increase metabolic activity (48, 49) and recent work indicates that even positive BOLD can reflect decreased metabolic activity in some contexts (50). iEEG responses provide greater clarity on these mechanisms as lipreading has been shown to elicit increased HGp in the posterior STG (15, 40), associated with increased neuronal firing rates (51), but beta suppression in the mid to anterior STG (15), associated with inhibitory processes (37). The notion that silent visual speech may suppress neural activity within portions of the auditory system has been theorized to reflect the optimized tuning of neurons specialized for auditory speech (52). Despite these differences in activation, above-chance classification was observed throughout the STG and pSTS suggesting two distinct mechanisms through which visual information is used to modulate phoneme populations. In posterior activations, we hypothesize that silent visual speech selectively activates matching phonetically or phonemically-tuned neurons in a categorical manner. Conversely, visemes may suppress activity in the STG in a targeted manner to inhibit incorrect representations in phonetically tuned neuronal populations (40) to indirectly refine the representation of correct phonetic features. While speculative, one possible explanation for why visemes avoid directly activating matching phonetically tuned neurons is to limit the potential for crossmodal hallucinations (53); i.e., hearing speech during silent lipreading.

Our current study demonstrates that visual speech information can be decoded in the auditory region, aligning with a growing body of research, including the findings of Brohl et al. (54), which underscores the multisensory nature of speech processing. While both studies underscore auditory cortex’s role in integrating visual speech, our approach, utilizing a combination of fMRI and intracranial EEG, provides unique spatial precision. This level of detail complements the temporal resolution offered by MEG, as used by Brohl et al., enabling a more comprehensive understanding of how visual and auditory speech information is integrated at different neural levels. Our controlled study design, focusing on specific parameters like stimulus length and voice onset time (VOT), offers distinct advantages over naturalistic designs such as those used by Brohl et al. (54). By minimizing confounds due to natural correlations between auditory and visual speech elements, we can more accurately isolate and test the encoding of visual speech information in the auditory cortex. This approach is particularly relevant when considering the findings of Brohl et al., who highlighted the auditory cortex’s role in reflecting unheard acoustic features during lip reading and established a behavioral relevance to these neural representations.

A limitation of the present work is that the small set of phonemes and visemes presented provide a limited account of the full distribution of phonemes and visemes present in English. This limitation was necessary to ensure adequate signal-to-noise ratios to enable classification of the individual phonemes and visemes, but future research can examine the full distribution of phoneme and viseme representations using more natural speech stimuli (55) in auditory-visual contexts using high-density intracranial electrodes. Data from such experiments would be predicted to show that phoneme tuning functions (the spatial selectivity of responses to a specific phoneme) will be more precise (narrower and more distinct from other phonemes) during auditory-visual speech compared to auditory-only speech. Moreover, we predict that phoneme and viseme spatial maps will imperfectly overlap (as the same viseme could denote ‘pet’ or ‘bet’) and that the dissimilarity in phoneme and viseme maps explain categorical shifts in perception during the McGurk effect (a perceptual illusion in which visual speech alters which phoneme is heard (56)).

In sum, the present studies provide the first direct evidence that the visemic features from lipread words are represented in the auditory system at a perceptual stage through spatially distributed neural activity. The data are consistent with the model that visemes share auditory representations with phonemes, which may serve to refine phonetic and phonemic population responses, to in turn support speech perception fluency. Importantly, while viseme identities could be classified from a majority of vertices in the STG that responded in general to visual speech, auditory regions likely encode a variety of other visual features (e.g., visual motion timing, visual parsing of speech rate, visual-derived spectral information) useful for facilitating speech perception processes in the natural environment.

## Methods

### fMRI Experiment

Planned analyses and sample size stopping justification for the fMRI study were pre-registered at OSF (https://osf.io/6fzwd/?view_only=60484583a2bb4dcdb8e27788c7c4c373). Minor deviations from the pre-registered protocol are noted throughout the methods section. The study was approved by the Institutional Review Board (IRB) of the University of Michigan.

### Subjects

FMRI data was acquired from *n* = 64 subjects (F = 47, M = 17) recruited from the University of Michigan’s Psychology paid-subject pool (individuals who had previously expressed interest in research studies) and through word of mouth. Subjects’ ages ranged from 18-32 (Mean: 22.87, SD = 3.29) and included 56 right-handed, 7 left-handed, and 1 ambidextrous individual. Written consent was obtained from each subject. subjects were paid USD $20 per hour for their time. Data was collected from each subject in a single session lasting approximately 1 hour and 15 minutes. Because power analyses using multivariate pattern analyses (MVPA) remain a challenge, we determined our sample size based on univariate power analysis (on the assumption that this would yield a minimum acceptable sample size). Sample size to detect visual-only effects was determined using data from the auditory-only condition in a preliminary sample using NeuroPower (using random field theory, cluster threshold p=.05, alpha=.05, n=27). Estimated sample sizes ranged from n=62 to 64 across the pairwise phoneme comparisons (/fafa/ vs /mama/, /fafa/ vs /kaka/, and /kaka/ vs /mama/) and n=64 was selected to ensure adequate power. No data from the visual-only condition was analyzed prior to submission of the pre-registration.

### Tasks, Stimuli and Experimental Design

We used an auditory and visual speech paradigm optimized for an event-related fMRI design. On each trial, subjects were presented with a three-alternative forced-choice task that consisted of either an auditory-only stimulus or a visual-only stimulus. Three types of phonemes; /fafa/, /kaka/ and /mama/ and three types of visemes; /fafa/, /kaka/ and /mama/ were used for this task and randomly mixed within blocks. These specific phonemes were chosen to maximize the differentiability between the individual phonemic representations in the neuronal populations of the STG (28, 44).

Fig. 1 shows the timing and structure of the task. Each trial for both the auditory-only and visual-only conditions lasted for 2 seconds. The auditory-only trials began with a fixation cross against a black screen, with the phonemes presented 250 ms after the appearance of the fixation cross. The visual-only trial began with the appearance of a female actor’s face on the screen, with lip movements beginning 250 ms after face onset. After the presentation of each auditory-only or visual-only trial, subjects were presented with 3 options (/fa/, /ka/, and /ma/) and were instructed to press one of three associated buttons on an MRI-safe button response box.

The first 24 subjects were shown response choices that always appeared in the same order (/fa/, /ka/, or /ma/) with a stable mapping between response choice and button (the index finger was always used to make the response for /fa/, the middle finger for /ka/ and the ring finger for /ma/). While performing the sample size estimates for our power analysis, we saw that the stable mapping between response choices and button presses resulted in response type differentiability in the motor cortex consistent with prior evidence for motor regions encoding information about finger movements (57). Hence, to counteract this effect and to negate the confounds of motor region responses during speech perception (58), we altered the pre-registered protocol for the remaining 40 subjects, who were shown response choices that were randomized after each trial. Group-level comparisons of MVPA data between these two sub-groups revealed significant differences only in the hand area of left M1/S1 for both the auditory-only and visual-only conditions.

Subjects had 1.25 seconds to respond to the answer choices. If the subject failed to register a response within 1.25 seconds, the trial was recorded as a missed response. Every trial was followed by a 5-6 second jitter period (sampled from a uniform random distribution) which acted as the intertrial interval (ITI) (59). In each run, subjects completed 60 trials that were split between 30 auditory-only and 30 visual-only trials, with 10 trials each for every phoneme and viseme; trial types and stimuli were randomly intermixed in each run.

In total, subjects completed five runs, resulting in 300 trials in total (150 phonemes, 150 visemes) during the task, with each run lasting 8 minutes and 30 seconds. Psychtoolbox was used for stimulus delivery and recording timing information and subject responses. Auditory stimuli were presented using fMRI compatible Avotec headphones that had integrated earmuffs in order to achieve maximum reduction of scanner noise. The sound level of stimuli was held constant for all subjects. While presenting auditory speech stimuli in an MRI scanner can be challenging, the undegraded nature of the auditory stimuli enabled near perfect accuracy throughout the task. A mirror system reflected the visual stimuli from an LCD projector onto a mirror (width of the mirror: 12cm, approximate viewing distance between eye and mirror: 15cm; width and height of the face on screen: 9cm x 12cm) located inside the magnet bore of the scanner.

### Data Exclusion Criteria

To ensure that subjects included in analyses demonstrated persistent attention throughout the task, we pre-registered exclusion criteria to remove subjects with behavioral accuracy rates less than 75% for either auditory-only or visual-only conditions: no subjects were excluded based on this cutoff.

### fMRI Data Collection

Subjects were scanned in a GE Discovery MR750 3.0 Tesla scanner with a Nova 32 channel standard adult-sized coil (Milwaukee, WI). One high-resolution T1-weighted structural image was obtained for each subject that was used in preprocessing, flip angle = 8, FOV = 25.6 mm, slice thickness = 1 mm, 256 slices. Then, for each of the five runs, functional T2*-weighted BOLD images were obtained using a multiband gradient-echo, echo planar imaging sequence with a resolution of 2.4 x 2.4 x 2.4 mm^3^, TR of 800 ms and, TE of 30 ms, Flip Angle of 52, for a total of 644 3D volumes of the whole brain with a FOV of 216 mm. To account for signal saturation, the task did not start until the first 10 TRs were acquired and discarded by the scanner in each run.

### Data Processing

fMRI data was reconstructed with realignment and fieldmap correction applied using SPM12 to each of the five T2* runs for inhomogeneity recovery of signal in the B0 field. Physiological noise was removed using RETROICOR (60). For both the univariate and multivariate analysis, preprocessing steps were completed using SPM12 (Wellcome Department of Cognitive Neurology, London, UK; https://www.fil.ion.ucl.ac.uk/spm/software/spm12/). We utilized The Decoding Toolbox (https://sites.google.com/site/tdtdecodingtoolbox/; version 3.997) for the whole-brain multivariate analyses.

### Preprocessing

Before preprocessing the functional images, SPM’s display tool was used to set the origin of the anatomical volumes for each subject manually by picking the location of the anterior commissure. After this, functional volumes were reconstructed and realigned, physiological noise was removed, and field map correction was applied. This was followed by slice time correction to account for acquisition time differences between slices for each of the whole brain functional volumes. This data was then co-registered to the subject’s anatomical space using a 4th degree B-spline, followed by segmentation of the tissues from the anatomical image with a forward deformation field. Information generated during the segmentation process was then used to transform the co-registered functional volumes into the standard MNI anatomical space with isotropic voxel volume dimensions of 2mm. The normalized data was then spatially smoothed using a full-width half maximum (FWHM) kernel of 5mm.

### Univariate Analyses

We performed a univariate, contrast-based analysis of auditory-only phonemes (averaged across the 3 phonemes) and visual-only visemes (averaged across the 3 visemes) in order to identify the regions that demonstrate significantly different activation patterns across stimulus types. We utilized a canonical hemodynamic response function with event duration set to 2 seconds for each of the phonemes (AuditoryFA + AuditoryKA + AuditoryMA) and visemes VisualFA + VisualKA + VisualMA) and 5.5 seconds for the fixation periods (Fixation). Event onsets times were defined as the moment when the fixation cross (for auditory trials) or face (visual trials) appeared on the screen.

In the first level analysis, whole brain beta maps were generated individually for all seven conditions for each of the 64 subjects. These maps also included information from regressors for motion correction (six head movement parameters). Freesurfer’s group-analysis pipeline was used for second level analyses (61). Specifically, each subject’s data was projected onto the cortical surface of the fsaverage subject (using the command mris_preproc) and smoothed using a FWHM of 10mm (using the command mri_surf2surf). General linear models were estimated with the command mri_glmfit separately for each hemisphere and condition, excluding motor and frontal areas due to the initial *n* = 24 subjects with consistent phoneme-motor mappings (analyses inclusive of all regions are shown in Supp. Fig. 1). Significant vertices were identified at the group level using the command mri_glmfit-sim using a vertex level threshold of *p* < .001 and cluster-level threshold of *p* < .05 (estimated with 10000 permutations) to control for multiple comparisons; p-values were adjusted for separate tests of the two hemispheres. All tests were two-tailed.

Univariate time-series were calculated from the individual subject volumes following fMRI realignment, unwarping with field map correction applied, and slice-timing correction. Volume data were epoched relative to phoneme or viseme onset for each individual trial, and the fourth volume (2.4 seconds post-stimulus) was used as a baseline for each trial. To quantify the relative differences in timing between phonemes and visemes, data were additionally normalized so that data in each condition for each subject ranged from 0-1.

### Multivariate Analyses

To identify regions that reliably differentiated classes of phonemes and classes of visemes, we performed searchlight based MVPA analyses. Preprocessing steps for univariate and multivariate analyses were matched except for the normalization and smoothing, such that for the multivariate analysis, these two steps were performed after the first level analysis was completed. For the decoding analysis, we utilized The Decoding Toolbox (62) with a LIBSVM (63) based support vector machine (SVM) implementation. For each of the individual subjects, we built a SVM classifier with a cross-validation scheme for the five runs. We used these classifiers to build two separate models: one to classify between the three phonemes and the other to classify between the three visemes. The phoneme models were constructed to identify voxels that reliably decoded the identity of each of the three phonemes while the viseme models were built to identify voxels that reliably decoded the identity for each of the three visemes. These models were implemented as independent whole-brain searchlight analysis in the first level of the MVPA model. For each of the models, beta estimates were calculated and extracted from a 3-voxel radius sphere. 4 fMRI runs were used for training and 1 run for testing in an iterative manner. The searchlight center was shifted through voxel-wise patterns throughout the brain to extract whole-brain accuracy maps for auditory-only and visual-only conditions. Chance-level accuracy (33.3%) was subtracted from individual subjects and conditions so that null-hypothesis values could be set to zero. Group-level analyses and multiple comparison corrections were performed using Freesurfer and matched those in the Univariate Analyses.

### ROI-Based Decoding Analyses

Following the whole-brain searchlight analysis, we selected five regions of interests from each hemisphere (ROI) based on results from literature (24, 43, 44). Four ROIs (STG, pSTS, fusiform, and hMT+) were pre-registered. The fifth ROI (V1/V2) was included in the classification analyses given the strong univariate response in the visual-only condition. ROIs were identified at the individual subject level based on Freesurfer aparc-aseg labeling (64). Selected labels included ‘superiortemporal’, ‘bankssts’,‘MT_exvivo.thresh’, the combinated labels ‘FG1.mpm.vpnl’ to ‘FG4.mpm.vpnl’, and the combined labels ‘V1_exvivo.thresh’ and ‘V2_exvivo.thresh’. Contrast beta estimates (condition vs fixation) were extracted for each subject, stimulus (6 phonemes and visemes), block, and ROI. SVM analyses were performed at the individual subject level with models trained on n-1 blocks (leave-one-out classification) using the ‘fitcecoc’ function in MATLAB.

### IEEG Experiment

The study was approved by the Institutional Review Boards (IRB) at the University of Michigan and Henry Ford Hospitals.

### Subjects and Recordings

*N* = 6 patients (2 female, 4 male) undergoing clinical evaluation using iEEG for intractable epilepsy consented to participate in this study under an institutional review board (IRB) approved protocol at the University of Michigan or Henry Ford hospital. Patients’ ages ranged from 12-39 years of age (mean = 29.7, std = 9.8) and 5 were right-handed (one patient self-reported to be ambidextrous). All patients were native English speakers. Clinically implanted depth electrodes (5 mm center-to-center spacing) and/or subdural electrodes (10 mm center-to-center spacing) were used to acquire iEEG data from subjects. IEEG data from a total of 459 electrodes were recorded from the six subjects. The type and location of electrodes implanted were based on the clinical needs of the patients. Electrodes were implanted within left auditory areas for 2 patients and right auditory areas for 4 patients. IEEG recordings were acquired at either 4096 Hz (*n* = 4 patients) or 1000 Hz (*n* = 2 patients) due to differences in clinical amplifiers.

### MRI and CT Acquisition and Processing

Preoperative T1-weighted magnetic resonance imaging (MRI) and postoperative computer tomography (CT) scans were acquired for all subjects. The preoperative T1 MRI was registered to the postoperative CT using SPM12 using the ‘mutual information’ method (65). The CT was not resliced or resampled. The localization of each electrode was performed using custom software (66). The algorithm works by identifying and segmenting electrodes from the CT image based on gray scale intensity, and projects subdural electrodes to the dura surface using the shape of the electrode disk to counteract post-operative compression. For all subsequent analyses including reconstruction of cortical surfaces, volume segmentation and anatomical labeling, the Freesurfer image analysis suite was utilized (http://surfer.nmr.mgh.harvard.edu/ (67, 68)).

### Task and Stimuli

Subjects were tested at their bedside in an Epilepsy Monitoring Unit using a laptop running Psychtoolbox (69). The task paradigm was adapted from a prior study (70) which was designed to behaviorally study multiple aspects of auditory-visual speech integration. The stimuli consisted of a female speaker who produced 40 commonly used 1-2 syllable words that each started with one of the four consonants: *‘b’, ‘f’, ‘g’, ‘d’* (10 of each). The phoneme in the second position of each of these words was generally balanced across each of the four groups. Each stimulus was recorded at a frame rate of 29.97 frames per second, and trimmed to 1100 ms in length. Further adjustments were made such that the first consonantal burst of each word occurred at 500 ms during the video playback by removing leading video frames.

Each subject underwent two task variants using the same stimuli and task design to increase trial numbers, and to reduce classifier overfitting. Supp. Fig. 6 shows the task schematic for both variants of the task. In variant one, subjects were presented with words one at a time, in one of two main conditions: auditory-only or visual-only. Subjects then identified the initial speech sound of the presented stimulus using a button press to select one of four options shown on the computer screen. For example, on a trial with the word *“bag”*, the options presented to the subject were *‘b’, ‘g’, ‘d’, ‘th’*. The paradigm included 40 trials per consonant in each main condition, such that each of the 40 words were presented 4 times in the visual-only condition and another 4 times in the auditory-only condition. This resulted in a total of 320 trials for each subject using task variant 1. The words used in our task are presented in Supplemental Table 6.

In task variant 2, subjects were presented with trials in one of four main conditions: auditory-only, visual-only, congruent audiovisual, or incongruent audiovisual. Task stimuli and instructions were the same as in variant 1. Variant 2 included 20 trials per consonant in each main condition. A second factor that was manipulated in this variant was the background noise level of the stimuli such that half of the words used in each condition were presented in either a low noise or a high noise context. In the low noise context, the auditory stimuli were presented as they were recorded (SNR = 32.2 dB SPL). In the high noise context, pink noise was added to reduce the signal-to-noise (SNR) ratio of the signals to −6 db SPL. Data were combined across the low and high noise contexts to increase the number of trials available for classification analyses. Only data from the auditory-only and visual-only conditions were included in analyses because they matched the main conditions obtained from Task variant 1 and as the auditory-visual conditions contained too few trials to be used in the classification analyses.

A total of 480 trials (Task variant 1: 320 trials, task variant 2: 160 trials) with 60 trials for each consonant (*‘b’, ‘g’, ‘d’, ‘f’*) per condition was obtained from the combined data of both task variants. Each subject received a randomized trial order. For the auditory-only condition, a gray rectangle was presented 500 ms before sound onset. Stimuli offset occurred 600 ms after sound onset time. In the visual-only condition, face onset occurred 500 ms before the time when phoneme onset would naturally occur. A wait time of 1.25 seconds was provided for the subjects to respond to each of the stimuli.

### IEEG Data Preprocessing

Data were preprocessed using bipolar referencing, such that signals from adjacent electrodes were subtracted in a pairwise manner. This ensured that the final signals of interest were obtained from neuronal populations that provided maximal localized responses (71). Analyses in auditory regions were restricted to electrodes (registered in MNI space) that were within 10 mm of the Freesurfer anatomical labels ‘superiotemporal’, ‘middletemporal’ or ‘supramarginal’. Excessively noisy electrodes were removed either manually or statistically by identifying electrodes with raw signals that were 5 SD greater in comparison to all other electrodes. For complementary analyses in visual regions, electrode locations were anatomically restricted to the ‘inferiortemporal’ and ‘fusiform’ labels.

Drift was removed from each channel (using residuals from fits to a 3rd order polynomial and high-pass filtering at 0.1 Hz). Power-line interference was removed by notch-filtering at 60 Hz and its harmonics. ERPs were extracted from this minimally processed signal. HGp activity was extracted from the continuous time-series after wavelet convolution and power transformation (70-150 Hz in 5 Hz intervals, wavelet cycles = 20 at 70 Hz, and increased linearly to maintain the same wavelet duration across frequencies). ERP and HGp data were segmented into 2 second epochs centered around speech onset time for a specific stimulus: trial onset was defined as the point when the initial consonant burst occurred. All data were then resampled to 1000 Hz.

Electrodes from both the left and right hemispheres were projected into the left hemisphere for analyses and visualization. This projection was performed by registering each subject’s skull-stripped brain to the Freesurfer cvs_avg35_inMN152 template image through affine registration using the Freesurfer function ‘mri_robust_register’ (72). Right hemisphere electrode coordinates were then reflected onto the left hemisphere across the sagittal axis.

### Classifiers for Calculating Decoding Accuracy

A support vector machine (73) classifier was utilized for calculating decoding accuracy. Classifiers for stimulus trials were built for individual subjects and group-level analyses were performed by combining results from individual subjects (subject as a random effect). Classification was performed on downsampled data (10 Hz except where stated otherwise) to reduce dimensional complexity. Phonemes and visemes were classified using a 4-fold multiclass classifier from 0 to 500 ms following sound onset time (or the corresponding point in the visual movie); electrodes and time-points were treated as dimensions in the classification of individual trials. Time-series analyses were performed independently at each time-point (1000 Hz) and accuracies were smoothed at the individual subject-level across 50 time points using the Matlab function ‘movmean’. Electrode-level analyses were performed on individual electrodes located in auditory regions.

### Similarity Analysis

To test whether auditory cortex showed a similar representation for phonemic and visemic information, we examined the similarity of the phoneme and viseme confusion matrices. Specifically, we paired each of the 16 cells in the two confusion matrices and used Pearson correlation to examine their relationship. Significance was calculated by randomly permuting the stimulus labels of each trial and repeating the full classification analysis *n* = 1000 times.

### Calculating Individual Subject Classification Significance

The four classes tested within each condition yielded chance levels of classification at 25%. To calculate significance above this chance level, we used binomial statistics for within-subject significance testing (74, 75). We used the ‘binocdf’ function in MATLAB for this, by considering two parameters: the number of trials, and probability of success at each instance (25%). This gives rise to a binomial chance-level probability that varies depending on the number of data points used for classification in each of the models that were built. This resulted in a chance probability of 29.58% (*p* = 0.05) for a 4-class classifier with 240 trials.

## Data Availability Statement

The data that support the findings of this study will be made openly available through the University of Michigan Deep Blue Repository.

## Acknowledgements

This study was supported by NIH Grants R00DC013828, 1R01DC020717, and R01NS094399.

