## Supplemental Materials for "Auditory cortex encodes lipreading information through spatially distributed activity"

**Supplemental Figure 1. Univariate activations without a frontal mask.** (a) phonemes vs fixation (b) visemes vs fixation, and (c) phonemes vs visemes. Phonemes evoked maximally increased activity in the STG bilaterally. Visemes evoked increased activity within bilateral visual cortex, left pSTS, right MT, and the STG bilaterally, though significantly weaker within the STG compared to phonemes. Colored regions reflect significant increases (red and yellow) or decreases (blues) in task-related activation (thresholded at p<.001 and corrected for multiple comparisons using cluster-statistics).

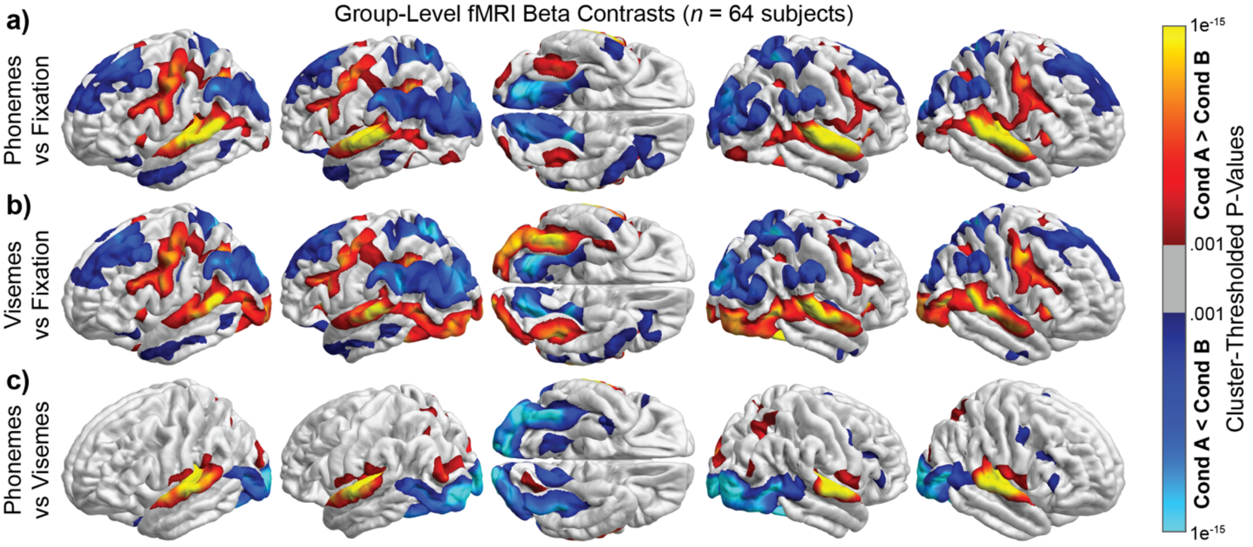

**Supplemental Figure 2.** Boxplots showing univariate group-level analyses of phonemes vs fixation (red) and visemes vs fixation (blue). Relative to fixation, phonemes evoked maximally increased activity in the STG bilaterally. Relative to fixation, visemes evoked increased activity within visual cortex, pSTS, hMT+, and the fusiform gyrus, along with suppression within the STG (including Heschl's gyrus). *p<.05, **p<.01, ***p<.001.

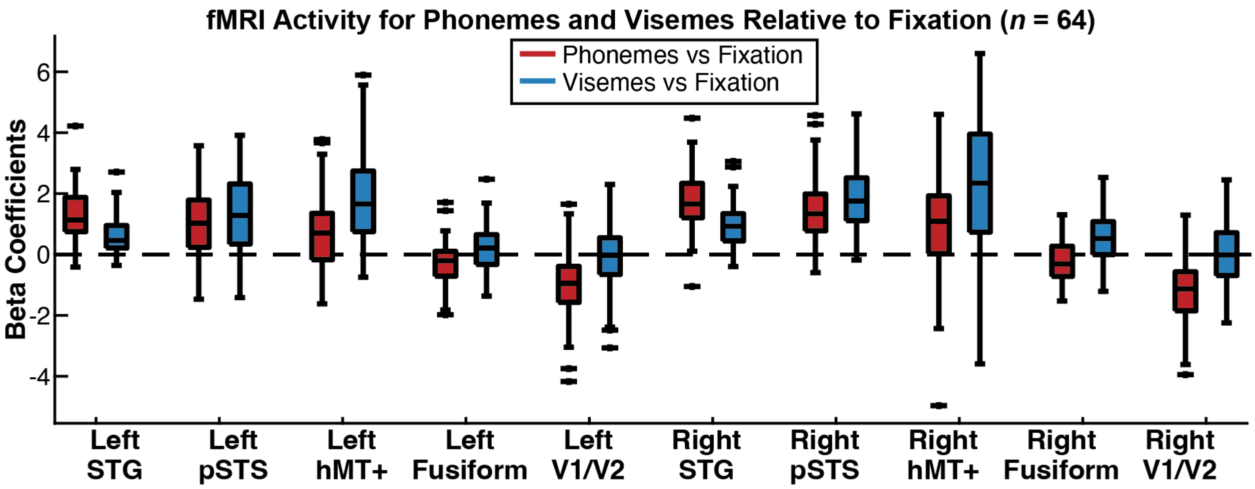

**Supplemental Figure 3.** BOLD time-series extracted from four regions (averaged across left and right hemispheres). Shaded region reflects 95% confidence intervals and gray boxes significantly different responses across the conditions corrected for multiple comparisons. Time-point zero reflects audio or movie onset.

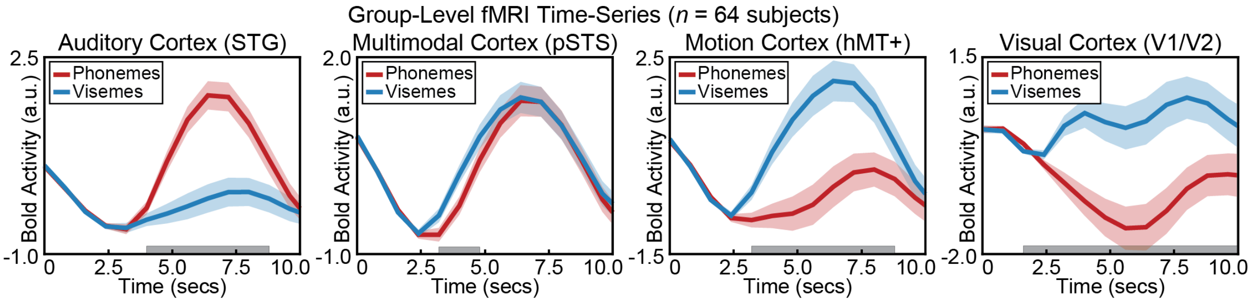

**Supplemental Figure 4.** Results of classification at all ROIs. Phonemes were significantly classified from the left STG and pSTS. Center line reflects the mean, colored box SE, and the tails 95% confidence intervals. *p<.05, **p<.01, ***p<.001. Chance accuracy is .333.

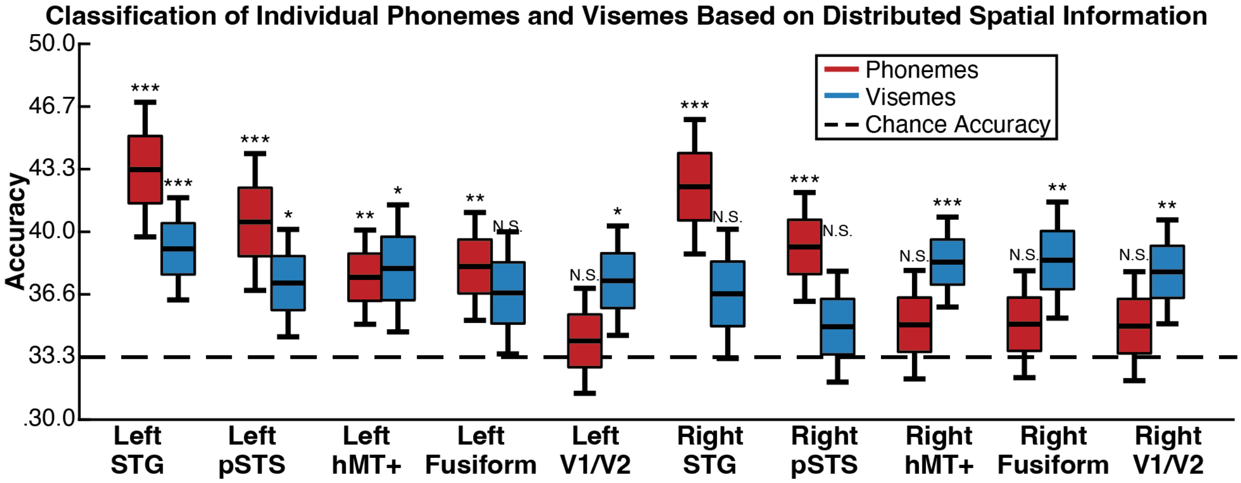

**Supplemental Figure 5.** Event-related spectral perturbations (ERSP) plots from electrodes within the anterior STG (top row), middle STG (middle row), and posterior STG (bottom row), averaged across subjects.

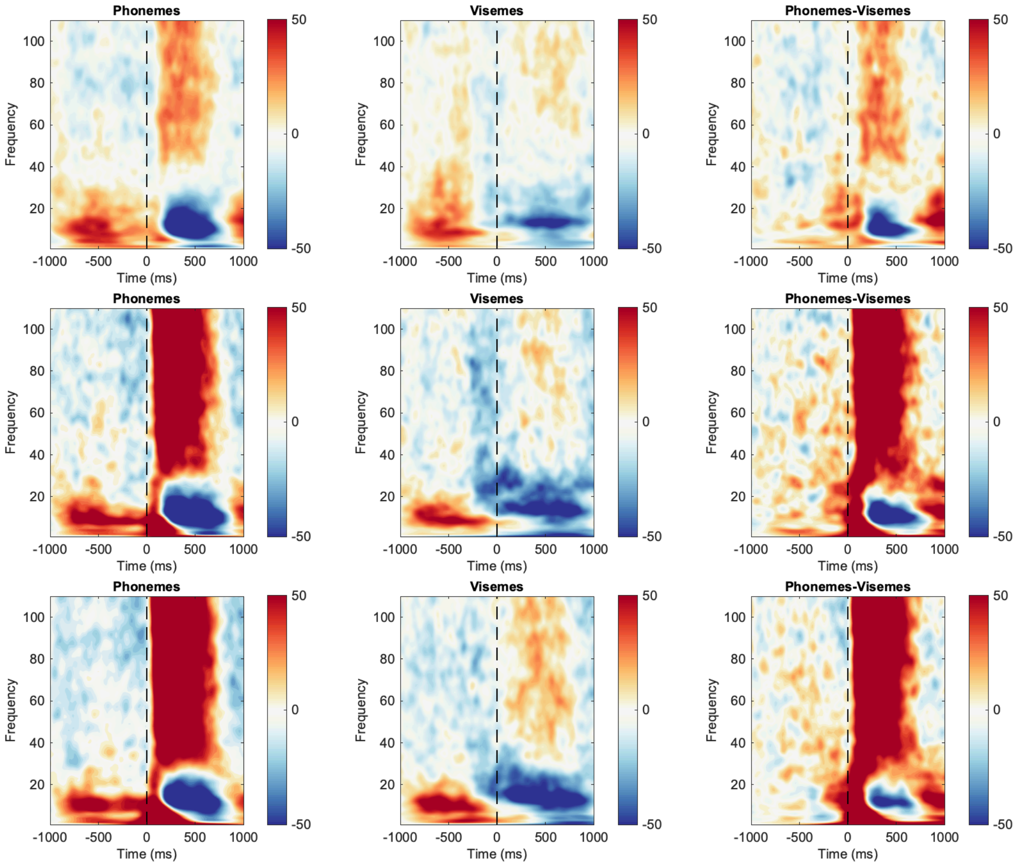

**Supplemental Figure 6.** Schematic of the iEEG task. All trials had an initial fixation period with a blank screen. In the auditory-only condition, a gray rectangle appeared on screen at 500 ms before onset of phonemes, which occurred at 0 ms. In the visual-only condition, following an initial fixation period, a face appeared on screen at 500 ms before the time when speech sounds would naturally begin. Stimuli offsets for both the auditory-only and visual-only conditions occurred at 600 ms after phoneme onset times. This was followed by a response window of 1250 ms in both conditions.

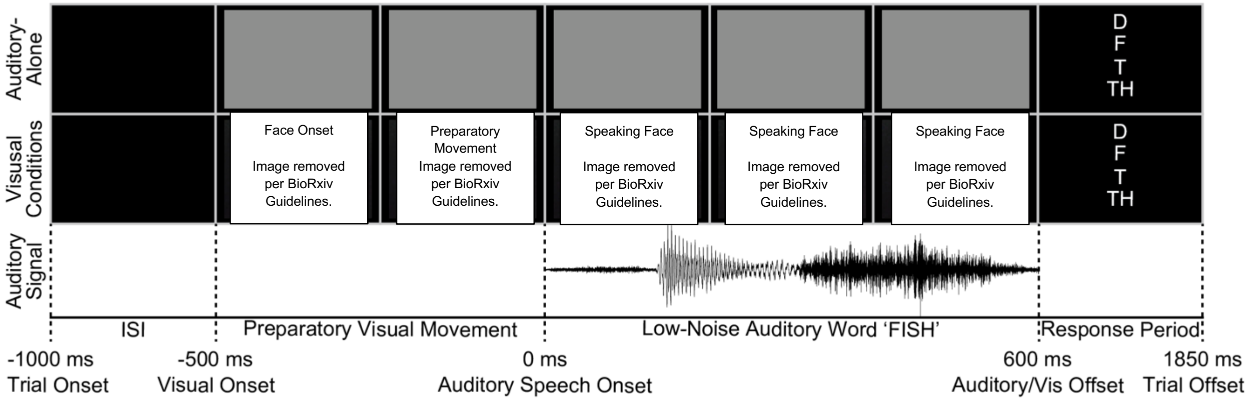

**Table 1**. MNI coordinates, statistics, and cluster sizes in the Univariate fMRI analysis for the **auditory-alone** condition. Anatomical regions involved are ordered by percentage of a structure that is encompassed by the cluster; if more than one region was included in a structure, only those with greater than 10% coverage of the anatomical region are reported. Negative log (p) values reflect areas with decreased activity relative to fixation, whereas positive log (p) values reflect areas with increased activity relative to fixation.

**Left**

| Cluster No. | Main Anatomical Regions Involved | Peak voxel coordinate in MNI | | | Cluster Number of Vertices | Cluster p-value | Min p (log) | ClusterSize (mm^2) |
|---|---|---|---|---|---|---|---|---|
| | | X | Y | Z | | | | |
| 1 | Cuneus (89.0%)<br>Lingual (87.7%)<br>Parahippocampal (79.5%)<br>Inferiorparietal (51.7%)<br>Pericalcarine (51.3%)<br>Lateraloccipital (32.6%)<br>Fusiform (25.9%)<br>Supramarginal (25.8%)<br>Precuneus (21.4%)<br>Superiorparietal (16.8%) | -23.5 | -38.2 | -13.1 | 20490 | 0.0002 | -15.7121 | 12126.35 |
| 2 | Transversetemporal (100.0%)<br>Superiortemporal (73.2%)<br>Bankssts (67.4%)<br>Supramarginal (22.6%) | -65.1 | -31.7 | 7.1 | 11179 | 0.0002 | 19.6639 | 5189.37 |
| 3 | Precuneus (21.5%)<br>Superiorparietal (20.1%) | -17.6 | -49.9 | 60.4 | 3672 | 0.0002 | -12.5229 | 1572.21 |
| 4 | Superiorparietal (22.2%) | -25.9 | -55.9 | 45 | 3405 | 0.0002 | 12.5911 | 1367.79 |
| 5 | Temporalpole (34.9%)<br>Middletemporal (16.9%) | -52.1 | 0.4 | -31.8 | 1388 | 0.0002 | -6.9865 | 1039.7 |
| 6 | Fusiform (23.9%)<br>Inferiortemporal (9.8%) | -41.8 | -47.3 | -14 | 1561 | 0.0006 | 7.3316 | 889.96 |
| 7 | Middletemporal (11.8%) | -57.3 | -49 | -5.2 | 683 | 0.0048 | -5.4388 | 410.25 |
| 8 | Lateraloccipital (7.1%) | -30.4 | -91.8 | -14 | 452 | 0.00918 | 6.2996 | 323.47 |

**Right**

| Cluster No. | Main Anatomical Regions Involved | Peak voxel coordinate in MNI | | | Cluster Number of Vertices | Cluster p-value | Min p (log) | ClusterSize (mm^2) |
|---|---|---|---|---|---|---|---|---|
| | | X | Y | Z | | | | |
| 1 | Cuneus (100.0%)<br>Pericalcarine (82.0%)<br>Parahippocampal (79.1%)<br>Lingual (78.5%)<br>Lateraloccipital (33.9%)<br>Fusiform (23.6%)<br>Superiorparietal (16.5%)<br>Precuneus (16.4%)<br>Inferiorparietal (12.0%) | 35.5 | -46.3 | -8.4 | 14841 | 0.0002 | -15.6293 | 10009.09 |
| **2** | Transversetemporal (100.0%)<br>Bankssts (88.8%)<br>Superiortemporal (77.0%)<br>Supramarginal (19.0%)<br>Middletemporal (14.0%)<br>Inferiorparietal (13.3%) | 65.3 | -21.3 | 1.1 | 11997 | 0.0002 | 22.6254 | 5280.77 |
| **3** | Superiorparietal (29.5%)<br>Precuneus (12.1%) | 14.2 | -50.9 | 64.3 | 3987 | 0.0002 | -14.2358 | 1697.9 |
| 4 | Supramarginal (21.7%)<br>Inferiorparietal (16.5%) | 59.9 | -28.4 | 40.5 | 3365 | 0.0002 | -9.8388 | 1596.84 |
| 5 | Fusiform (32.1%) | 40.5 | -57.8 | -11.3 | 1795 | 0.0006 | 9.8941 | 1015.48 |

| | | | | | | | | |
|---|---|---|---|---|---|---|---|---|
| **6** | Superiorparietal (8.8%)<br>Inferiorparietal (5.6%) | 27.7 | -55.6 | 45.7 | 1443 | 0.003 | 7.1896 | 594.55 |
| **7** | Middletemporal (9.2%)<br>Inferiortemporal (6.2%) | 46.4 | -5.2 | -32.7 | 723 | 0.005 | -6.7801 | 468.13 |
| 8 | Lateraloccipital (9.1%) | 30.8 | -93.6 | -12.1 | 540 | 0.00579 | 6.1315 | 438.21 |

**Table 2**. MNI coordinates, statistics, and cluster sizes in the Univariate fMRI analysis for the **visual-alone** condition. Anatomical regions involved are ordered by percentage of a structure that is encompassed by the cluster; if more than one region was included in a structure, only those with greater than 10% coverage of the anatomical region are reported. Negative log (p) values reflect areas with decreased activity relative to fixation, whereas positive log (p) values reflect areas with increased activity relative to fixation.

**Left**

| Cluster No. | Main Anatomical Regions Involved | Peak voxel coordinate in MNI | | | Cluster Number of Vertices | Cluster p-value | Min p (log) | ClusterSize (mm^2) |
|---|---|---|---|---|---|---|---|---|
| | | X | Y | Z | | | | |
| 1 | Cuneus (63.6%)<br>Inferiorparietal (55.6%)<br>Precuneus (54.4%)<br>Superiorparietal (42.0%)<br>Supramarginal (26.5%)<br>Lateraloccipital (16.8%) | -50.2 | -50.3 | 32 | 17129 | 0.0002 | -13.9686 | 8914.83 |
| 2 | Bankssts (67.8%)<br>Lateraloccipital (55.1%)<br>Superiortemporal (53.9%)<br>Transversetemporal (53.1%)<br>Fusiform (41.2%)<br>Inferiortemporal (17.2%)<br>Supramarginal (15.6%)<br>Middletemporal (10.8%) | -64.1 | -38.4 | 10.8 | 14525 | 0.0002 | 15.7542 | 7960.44 |
| 3 | Parahippocampal (65.3%)<br>Lingual (54.2%)<br>Fusiform (21.8%)<br>Pericalcarine (18.1%) | -18.2 | -78 | -10.2 | 4923 | 0.0002 | -14.9722 | 2998.1 |
| 4 | Superiorparietal (20.9%) | -26.9 | -52.1 | 44.4 | 3233 | 0.0002 | 12.6031 | 1280.5 |
| 5 | Middletemporal (19.3%)<br>Temporalpole (13.3%) | -52.6 | -1.7 | -31.7 | 1374 | 0.0002 | -5.8522 | 1039.08 |
| 6 | Middletemporal (12.3%) | -58.6 | -49.2 | -4.5 | 692 | 0.0048 | -5.6698 | 411.73 |

**Right**

| Cluster No. | Main Anatomical Regions Involved | Peak voxel coordinate in MNI | | | Cluster Number of Vertices | Cluster p-value | Min p (log) | ClusterSize (mm^2) |
|---|---|---|---|---|---|---|---|---|
| | | X | Y | Z | | | | |
| 1 | Bankssts (94.4%)<br>Superiortemporal (62.4%)<br>Lateraloccipital (60.0%)<br>Fusiform (59.3%)<br>Transversetemporal (27.5%)<br>Inferiortemporal (21.2%)<br>Middletemporal (21.0%)<br>Inferiorparietal (17.0%)<br>Supramarginal (13.6%) | 62.6 | -36.2 | 9.5 | 17955 | 0.0002 | 20.597 | 9650.22 |
| **2** | Cuneus (59.9%)<br>Superiorparietal (54.8%)<br>Precuneus (37.1%)<br>Lateraloccipital (16.4%)<br>Inferiorparietal (11.5%) | 10.8 | -59.5 | 60.3 | 11634 | 0.0002 | -16.3323 | 6370.23 |
| **3** | Parahippocampal (61.8%)<br>Lingual (48.3%)<br>Pericalcarine (20.9%)<br>Fusiform (20.1%) | 35.8 | -45.5 | -8.8 | 4315 | 0.0002 | -16.8827 | 2832.76 |
| 4 | Supramarginal (21.2%)<br>Inferiorparietal (13.6%) | 59.9 | -28.8 | 38.5 | 3046 | 0.0002 | -8.0376 | 1447.59 |
| 5 | Superiorparietal (7.6%)<br>Inferiorparietal (5.3%) | 27.7 | -55 | 45.5 | 1293 | 0.0022 | 7.5652 | 529.37 |

| | | | | | | | | |
|---|---|---|---|---|---|---|---|---|
| **6** | Middletemporal (4.7%)<br>Inferiortemporal (4.3%) | 46 | -4.5 | -32.7 | 418 | 0.01296 | -5.3922 | 279.2 |

**Table 3**. MNI coordinates, statistics, and cluster sizes in the Univariate fMRI analysis for the **auditory-alone vs visual-alone** conditions. Anatomical regions involved are ordered by percentage of a structure that is encompassed by the cluster; if more than one region was included in a structure, only those with greater than 10% coverage of the anatomical region are reported. Negative log (p) values reflect areas with greater activity in visual-alone compared to auditory-alone conditions, whereas positive log (p) values reflect areas with increased activity in auditory-alone vs visual-alone conditions.

| **Left** | | | | | | | | |
|---|---|---|---|---|---|---|---|---|
| **Cluster No.** | **Main Anatomical Regions Involved** | **Peak voxel coordinate in MNI** | | | **Cluster Number of Vertices** | **Cluster p-value** | **Min p (log)** | **ClusterSize (mm^2)** |
| | | **X** | **Y** | **Z** | | | | |
| 1 | Lateraloccipital (73.2%)<br>Pericalcarine (69.2%)<br>Lingual (54.1%)<br>Fusiform (50.3%)<br>Parahippocampal (18.3%)<br>Cuneus (17.0%)<br>Bankssts (16.9%)<br>Middletemporal (12.6%)<br>Inferiortemporal (10.9%) | -14.4 | -100.2 | -3.5 | 13393 | 0.0002 | -19.5886 | 8411.5 |
| 2 | Transversetemporal (100.0%)<br>Superiortemporal (68.0%)<br>Supramarginal (16.0%) | -50 | -15.9 | 3.3 | 7434 | 0.0002 | 17.6538 | 3287.17 |
| 3 | Inferiorparietal (12.9%)<br>Lateraloccipital (8.5%)<br>Cuneus (6.6%)<br>Superiorparietal (6.5%) | -34.8 | -85.2 | 15.4 | 2338 | 0.0002 | 11.2267 | 1578.76 |
| 4 | Lingual (12.2%)<br>Fusiform (10.4%)<br>Parahippocampal (6.8%) | -30.7 | -59.4 | -7.6 | 1126 | 0.0012 | 6.2173 | 665.9 |
| 5 | Precuneus (16.8%)<br>Superiorparietal (2.3%) | -7.7 | -47.3 | 48.2 | 1472 | 0.0018 | 7.2845 | 620.65 |
| 6 | Superiorparietal (4.4%) | -17.3 | -64.7 | 46 | 455 | 0.0292 | 4.2978 | 209.91 |
| **Right** | | | | | | | | |
| **Cluster No.** | **Main Anatomical Regions Involved** | **Peak voxel coordinate in MNI** | | | **Cluster Number of Vertices** | **Cluster p-value** | **Min p (log)** | **ClusterSize (mm^2)** |
| | | **X** | **Y** | **Z** | | | | |
| 1 | Pericalcarine (88.4%)<br>Lateraloccipital (79.2%)<br>Lingual (65.4%)<br>Fusiform (53.6%)<br>Bankssts (49.5%)<br>Cuneus (28.6%)<br>Parahippocampal (21.4%)<br>Inferiortemporal (20.9%)<br>Middletemporal (20.4%)<br>Inferiorparietal (11.6%) | 40.6 | -59.7 | -16.6 | 16668 | 0.0002 | -19.1577 | 10482.03 |
| **2** | Transversetemporal (100.0%)<br>Superiortemporal (58.7%) | 44.3 | -21.5 | 6.3 | 5541 | 0.0002 | 20.5555 | 2388.74 |
| **3** | Superiorparietal (12.5%) | 20.6 | -64.6 | 40.6 | 1273 | 0.0014 | 6.9392 | 689.91 |
| 4 | Precuneus (21.6%) | 16.2 | -45.1 | 52.4 | 1767 | 0.0018 | 7.4801 | 635.39 |
| 5 | Inferiorparietal (6.3%) | 35.1 | -80.1 | 19 | 742 | 0.00619 | 7.0068 | 420.02 |
| **6** | Entorhinal (16.0%)<br>Fusiform (9.2%) | 37 | -5 | -35 | 642 | 0.00918 | -5.4481 | 362.01 |

**Table 4** full statistics for MVPA fMRI analysis for the auditory-alone condition, main anatomical regions involved are ordered by percentage of the cluster.

| Cluster No. | Main Anatomical Regions Involved | Peak voxel coordinate in MNI (x y z) | | | Cluster Number of Vertices | Cluster p-value | Max log (p) | ClusterSize (mm^2) |
|---|---|---|---|---|---|---|---|---|
| **Left Brain** | | | | | | | | |
| 1 | STS (89%)<br>STG (61.8%)<br>SMG (39.2%)<br>A1 (33.1%)<br>MT (8.1%) | -64.1 | -33.5 | 9.4 | 10855 | 0.0002 | 9.0701 | 5047.12 |
| 2 | SPL (27.3%)<br>IPL (15.8%)<br>SMG (8.1%) | -33.8 | -55.7 | 35.4 | 4791 | 0.0004 | 5.7955 | 1941.39 |
| 3 | IT (13.8%)<br>FFA (8.1%)<br>LOC (3.4%) | -47.1 | -58.6 | -7.5 | 1206 | 0.00719 | 5.2061 | 693.14 |
| **Right Brain** | | | | | | | | |
| 1 | A1 (57.4%)<br>STG (45%)<br>STS (26.7%)<br>SMG (0.1%) | 59.4 | -15.9 | 2.4 | 4134 | 0.0002 | 9.052 | 1841.88 |
| 2 | MT (14.2%)<br>STS (7.1%)<br>IPL (0.1%) | 56.5 | -59.6 | 3.7 | 887 | 0.01396 | 4.3151 | 490.85 |

**Table 5**. Full statistics for MVPA fMRI analysis for the visual-alone condition, main anatomical regions involved are ordered by percentage of the cluster.

| Cluster No. | Main Anatomical Regions Involved | Peak voxel coordinate in MNI (x y z) | | | Cluster Number of Vertices | Cluster p-value | Max log (p) | ClusterSize (mm^2) |
|---|---|---|---|---|---|---|---|---|
| **Left Brain** | | | | | | | | |
| 1 | STS (76.9%)<br>STG (42.6%)<br>SMG (29.9%)<br>A1 (3.9%)<br>MT (3.3%) | -58.4 | -26.1 | 25.3 | 7652 | 0.0002 | 5.8515 | 3572.86 |
| 2 | LOC (47.4%)<br>Lingual (15.1%)<br>V1 (11.9%)<br>FFA (4.2%)<br>IPL (<0.1%) | -19.5 | -99.8 | 2.3 | 4085 | 0.0002 | 6.0987 | 3288.8 |
| 3 | SMG (20.1%) | -52.9 | -40.8 | 45.8 | 1731 | 0.00639 | 4.5852 | 698.48 |
| 4 | SPL (8%)<br>IPL (3.8%) | -28.3 | -66.2 | 37.3 | 1132 | 0.01316 | 3.4307 | 502.88 |
| **Right Brain** | | | | | | | | |
| 1 | LOC (67.2%)<br>Lingual (20.1%)<br>V1 (10.3%)<br>FFA (6%)<br>IPL (0.3%) | 24.4 | -85.7 | -8.2 | 5291 | 0.0002 | 7.9634 | 4110.07 |
| 2 | STG (12.5%)<br>A1 (7.7%)' | 57.3 | -16.8 | 2.4 | 918 | 0.01931 | 4.1714 | 424.48 |

**Table 6.** Word stimuli used in the fMRI and ECoG experiments. Each word began with one of four consonants ('B', 'F', 'G', or 'D'). Among the 40 distinct words used, 10 words with different initial vowels were paired with the consonant. Vowels were matched across consonant categories. Each word was repeated 6 times within each condition.

| ***Consonant*** | ***Vowel 1*** | ***Vowel 2*** | ***Vowel 3*** | ***Vowel 4*** | ***Vowel 5*** | ***Vowel 6*** | ***Vowel 7*** | ***Vowel 8*** | ***Vowel 9*** | ***Vowel 10*** |
|---|---|---|---|---|---|---|---|---|---|---|
| ***B*** | Bag | Base | Beard | Bible | Bid | Bank | Beer | Bias | Bill | Bye |
| ***F*** | Fad | Fat | Fill | Fine | Fist | Fast | File | Fish | Fit | Five |
| ***G*** | Gag | Gas | Guild | Guile | Gig | Gang | Gear | Guise | Gill | Guy |
| ***D*** | Dad | Dash | Dill | Dine | Disk | Dance | Dial | Dish | Digs | Dive |

**Table 7.** Single-trial ERP classification accuracy (%) for individual subjects in the auditory-only and visual-only conditions.

| **Subject** | **Auditory-only** | **Visual-only** |
|---|---|---|
| 1 (UM1219) | 31.25 ($p = 0.012$) | 31.67 ($p = 0.008$) |
| 2 (UM1208) | 40.83 ($p < 0.001$) | 40.00 ($p < 0.001$) |
| 3 (HF1187) | 39.58 ($p < 0.001$) | 30.42 ($p < 0.001$) |
| 4 (UM1223) | 45.83 ($p < 0.001$) | 34.17 ($p < 0.001$) |
| 5 (HF1183) | 32.92 ($p = 0.002$) | 31.67 ($p = 0.008$) |
| 6 (UM1234) | 31.11 ($p = 0.006$) | 31.21 ($p = 0.005$) |
| Group | M = 36.92 %, SD = 6.06 % | M = 33.19 %, SD = 3.57 % |

**Table 8.** Group statistics of SVM decoding results of each one of the auditory-alone and visual-alone stimuli, compared to chance (of 25%).

| | Aud B | Aud F | Aud G | Aud D | Vis B | Vis F | Vis G | Vis D |
|---|---|---|---|---|---|---|---|---|
| **P Value** | 0.0483 | 0.0017 | 0.0617 | 0.0096 | 0.0288 | 0.0041 | 0.0579 | 0.0364 |
| **T stat** | 2.0425 | 5.2523 | 1.8514 | 3.4000 | 2.4559 | 4.2253 | 1.9007 | 2.2670 |
| **Degrees of Freedom** | 5 | 5 | 5 | 5 | 5 | 5 | 5 | 5 |
| **Standard Deviation** | 0.0930 | 0.1083 | 0.0987 | 0.0680 | 0.0886 | 0.0732 | 0.0710 | 0.0606 |
